# A tau class Glutathione-S-Transferase (OsGSTU5) acts as a negative regulator of VirE2 interaction with T-DNA during *Agrobacterium* infection in rice

**DOI:** 10.1101/2021.03.25.436940

**Authors:** Madhu Tiwari, Neelam Gautam, Yuvraj Indoliya, Maria Kidwai, Arun Kumar Mishra, Debasis Chakrabarty

## Abstract

During *Agrobacterium*-mediated transformation (AMT), T-DNA along with several virulence proteins like VirD2, VirE2, VirE3, VirD5, and VirF enter into the plant cytoplasm. VirE2 is supposed to serve as single-stranded DNA binding (SSB) protein and assist the cytoplasmic trafficking of T-DNA inside the host cell. In the present study, a rice glutathione-S-transferase (OsGSTU5) that interacts with VirE2 protein in plant cytoplasm has been identified. OsGSTU5 is observed to be involved in post-translational glutathionylation of VirE2 protein (gVirE2). *In silico* analysis revealed that ‘gVirE2+ssDNA’ complex is structurally less stable than ‘VirE2+ ssDNA’ complex. The gel shift activity confirms the attenuated SSB property of gVirE2 over VirE2 protein under *in vitro* condition. Moreover, knock-down and overexpression *OsGSTU5* phenotypes of rice showed increased and decreased T-DNA expression, respectively after *Agrobacterium* infection. The present finding convincingly establishes the role of OsGSTU5 as defense protein in rice that can further serve as an important target for modulation of AMT efficiency in rice.

## Introduction

*Agrobacterium tumefaciens* (AT) is a soil-borne plant pathogen that causes crown gall disease in plants (Chilton et al. 1977). The pathogenicity of *Agrobacterium* is governed by the specific plasmid termed as Tumor inducing (Ti) plasmid. During infection, a part of a Ti-plasmid i.e., T-DNA, is generated by VirD1-VirD2 endonuclease inside the bacteria (Herrera-Estrella et al. 1988; Wang et al. 1984; Yanofsky et al. 1986). T-DNA is transferred into the host cell and subsequently integrated to its genome. Native T-DNA contains oncogenes which when integrated to the host genome cause hormonal imbalance in host plants and result in gall formation (Chilton et al. 1977; Zupan et al. 2000). The native T-DNA can be replaced by any gene of interest and it can be utilized for the development of transgenic plants with desirable traits (Caplan et al. 1983; Fraley et al. 1983). This method is called *Agrobacterium*-mediated transformation (AMT) and is extensively utilized for development of various transgenic plants (Gelvin 2009; Valentine 2003).

*Agrobacterium* has remarkably wide host range, but majority of the monocot plants are not within the natural host range (Gelvin 2003). Interestingly, the current state of literature reveals that under laboratory conditions AMT is not only limited to its natural hosts (usually dicot plants) but also useful for transforming the non-hosts like, monocots (Cheng et al. 1997; Frame et al. 2002; Hiei et al. 1994; Reyes et al. 2010; Ziemienowicz et al. 2012), yeast (Bundock et al. 1995), fungi (De Groot et al. 1998; Li et al. 2017), and human cells (Lacroix and Citovsky 2018).

During AMT, in addition to T-DNA, several other virulence proteins viz. VirD2, VirE2, VirE3, VirD5, and VirF are also transferred to the host cell (Schrammeijer et al. 2003; Vergunst et al. 2005). T-DNA is transferred to the host cell as a single strand DNA (ssT-DNA) which is associated with VirD2 protein at 5’ end, while the other virulence proteins enter into the host cell independently via a type 4 secretion system (T4SS) (Li and Christie 2018; Vergunst et al. 2003; Vergunst et al. 2005). Among all the Vir encoded proteins, VirE2 is the most abundant protein of *Agrobacterium* (Engström et al. 1987), which utilizes the host endomembrane compartments for its transfer and trafficking into the host cell (Li and Pan 2017; Tu et al. 2018). VirE2 binds ssT-DNA without any sequence specificity and forms a solenoid structure (Abu-Arish et al. 2004; Christie et al. 1988; Citovsky et al. 1989; Dym et al. 2008; Sen et al. 1989). It is hypothesized that VirE2 protect ssT-DNA from host nucleolytic degradation and maintains T-DNA integrity in host cell (Howard and Citovsky, 1990, Yusibov et al. 1994, Rossi et al, 1996, Ziemienowicz et al. 2001). Later, Yang et al. (2017) showed that, host myosin XI-K–powered ER/actin network systems are involved in cytoplasmic trafficking of VirE2 and T-DNA (Yang et al. 2017). However, the mechanism of VirE2 mediated nuclear import of T-DNA is still unclear due to some contrasting reports. Some studies showed that cytoplasmic localization of VirE2 (Shi et al. 2014). On the contrary, other studies reported the delivery of VirE2 in the nucleus of plants under natural infection conditions (Li et al. 2014; Li et al. 2020). In Arabidopsis, the (VirE2 interacting protein 1) VIP1-VirE2 interaction was suggested to facilitate the nuclear import of VirE2 and thus T-DNA import too (Loyter et al. 2005; Tzfira et al. 2001). Later, VIP1 was identified as a MAP-Kinase activated protein which amplifies the defense response of the plant by activating the secondary defense related genes (Djamei et al. 2007, Pitzschke et al. 2009, Lacroix and Citovsky 2013).

Globally, VirE2 is found to protect T-DNA, assist its trafficking into the host cytoplasm by utilizing existing host cellular machinery and possibly involve in T-DNA nuclear import. Under laboratory conditions the AMT protocol has been established in monocot plants like rice (Hiei et al. 1997; Hiei et al. 1994), suggesting the meanwhile activation and participation of host proteins. Until now, VirE2 interacting protein has not been explored in any monocot plant. Keeping this gap in mind, the present investigation has been carried out to identify and functionally characterize the host proteins that play a key role by interacting VirE2 proteins during AMT in rice.

In this study, a cDNA library of *Agrobacterium* inoculated rice calli was used for identification of VirE2 interacting protein in rice using a yeast two-hybrid assay. Through reporter-based interaction, a tau class glutathione-S-transferase, OsGSTU5 (*Os09g20220*) from rice which interacts with the VirE2 protein, has been identified. The observed interaction was further confirmed by bimolecular fluorescence complementation (BiFC) and pull-down assays. Later, OsGSTU5 mediated in vitro glutathionylation activity revealed the post-translational modification of VirE2 into its glutathionylated form. In addition, the decrease ssT-DNA binding property of glutathionylated VirE2 (gVirE2) protein over non-glutathionylated VirE2 protein (VirE2) has also been observed. In order to characterize functionally the probable role of GSTU5 in rice, we prepared the knockdown and overexpression lines which respectively showed the higher and lower T-DNA expression. In brief, GSTU5 can be considered as a defense protein that affects the AMT efficiency in rice.

## Materials and methods

### Experimental material

In the present study, Nipponbare rice (*Oryza sativa* L. ssp. *japonica*) and *Agrobacterium tumefaciens* strain EHA101(Hood et al. 1986) have been used. For cloning DH5α and protein expression BL21 (DE3) (Invitrogen) strains of *E*.*coli* were used. Seeds of Nipponbare rice (*Oryza sativa* L. ssp. Japonica) variety were dehusked and washed with autoclaved MiliQ water (3 times). Seeds were surface sterilized in 70% ethanol for 90 sec followed by washing with MiliQ water (3 times). Seeds were washed using 0.1% HgCl_2_ disinfectant for 30 sec followed by thorough washing with MiliQ water (5-8 times). In addition to it, 2% sodium hypochlorite solution with a few drops of Tween20 was used to wash the seeds for 1 hour with gentle shaking at 15 min intervals. The process was followed by washing with MiliQ water in order to remove any trace of chemical. Next, sterilized rice seeds were placed on callus induction media (N6). After callus formation, the gene of interest was transformed into rice calli via *Agrobacterium*-mediated transformation.

### Plasmid constructs

*VirE2* (Gene ID: 6382154, NCBI reference protein YP_001967551.1, Locus tag pTiBo153) ORF was amplified from the EHA101 *Agrobacterium* strain (Hood et al. 1986), harbouring the pTiBo542 helper plasmid and cloned to pGBKT7 cloning vector (Clontech, Mountain View, USA) (Ramalingam et al. 2015). pGBKT7 was linearized with *BamHI* and *EcoRI* and fused with the PCR amplified VirE2 ORF (with VirE2 infusion primer). The primer sequence has been provided in Table S1.

Full-length OsGSTU5 was obtained from the rice genome annotation project (http://rice.plantbiology.msu.edu/cgi-bin/sequence_display.cgi?orf=LOC_Os09g20220.1) and cloned to the pGAD cloning vector (Clontech, Mountain View, USA) (Ramalingam et al. 2015) as an *EcoRI-BamHI* fragment. In this construct, *EcoRI* was taken as forward (F) and *BamHI* was taken as reverse primers (R). The primers sequence has been given in as GSTU5 (pGAD) primers in Table S1.

To trace co-expression of both genes in onion peel during the BiFC assay, *VirE2* and *OsGSTU5* ORFs were PCR amplified and cloned as *XhoI*(F)*-KpnI*(R) and *EcoRI*(F)*- BamHI*(R) fragments into pSAT1-nEYFP-C1(Citovsky et al. 2006) vector generating pSAT1-nEYFP-C1-*VirE2* and pSAT1-cEYFP-C1B-*GSTU5*, respectively. *OsGSTU30* (*Os10g38600*) was used as a negative control in the split YFP assay. *OsGSTU30* was PCR amplified as a *HindIII*(F)-*KpnI*(R) fragment and cloned to the pSAT1-cEYFP-C1B vector. All the primers used are termed as BiFC primer and they are listed in Table S1.

For subcellular localization study of *GSTU5*, the *OsGSTU5* ORF was amplified by PCR and fused as a *EcoRI*(F)*-BamHI*(R) fragment into the pSAT6-EYFP-C1 (Citovsky et al. 2006) vector generating the pSAT6-EYFP-C1-*GSTU5* construct. The same set of primer is used in localization study as used in BiFC assay earlier.

Both *VirE2* and rice *GSTU5* ORFs were amplified by PCR and cloned into pETSUMO vector (Invitrogen) (Wang et al. 2010) for protein expression. The primer sequence has been provided in Table S1.

To generate the Knockdown (KD) *GSTU5* construct, primers were synthesized using the WMD3, http://wmd3.weigelworld.org/cgi-bin/webapp.cgi website tool (ImiR-s, IImiR-a, IIImiR*s, IVmiR*a). Those primers were replaced with the *Oryza sativa* miRNA precursor, osa-miR158, previously cloned into the pNW55 vector (Addgene) (Chen et al. 2012). The original miRNA sequence was replaced by the *KDGSTU5* in the pNW55 vector via PCR-based site-directed mutagenesis. *KDGSTU5* and overexpression *GSTU5* constructs were PCR-amplified and fused as a *BamHI*(F)*-KpnI*(R) and *BamHI*(F)*- KpnI*(R) fragments respectively into the pIRS154 vector (a pCambia derivative, gifted by Emmanuel Guiderdoni, CIRAD, France) (Chen et al. 2008) under the control of the Ubiquitin maize promoter. All primer sequences are listed in Table S1.

### Library Preparation

Initially, the *Agrobacterium*-infected rice calli were screened for T-DNA (β-Glucuronidase; GUS) expression by using the GUS histochemical assay. The cDNA library was prepared from japonica rice calli. The rice calli were incubated with the *Agrobacterium* strain EHA101 harbouring a pIG121Hm^51^ vector. Here the *uidA* (*GUS*) gene was used as the T-DNA. GUS expression of rice calli was tested from 45 minutes-120 hours post-inoculation (Fig. 1a). Total mRNA was isolated from rice calli 24, 48, and 72 hours post-inoculation, respectively. Next, the cDNA library was prepared using the “mate and plate” library preparation system (Clontech, Mountain View, USA). The prepared library, thereafter, was directly transformed into the *Saccharomyces cerevisiae* Y187 yeast strain (*MATa, ura3-52, his3-200, ade2-101, trp1-901, leu2-3, 112, gal4*Δ, *gal80*Δ, *met–, URA3: GAL1UAS–Gal1TATA–LacZ MEL1*) via the Yeastmaker™ Yeast Transformation System 2 kit (Clontech, Mountain View, USA). An agarose gel image of the cDNA library is shown in Fig. 1b.

**Fig 1.**
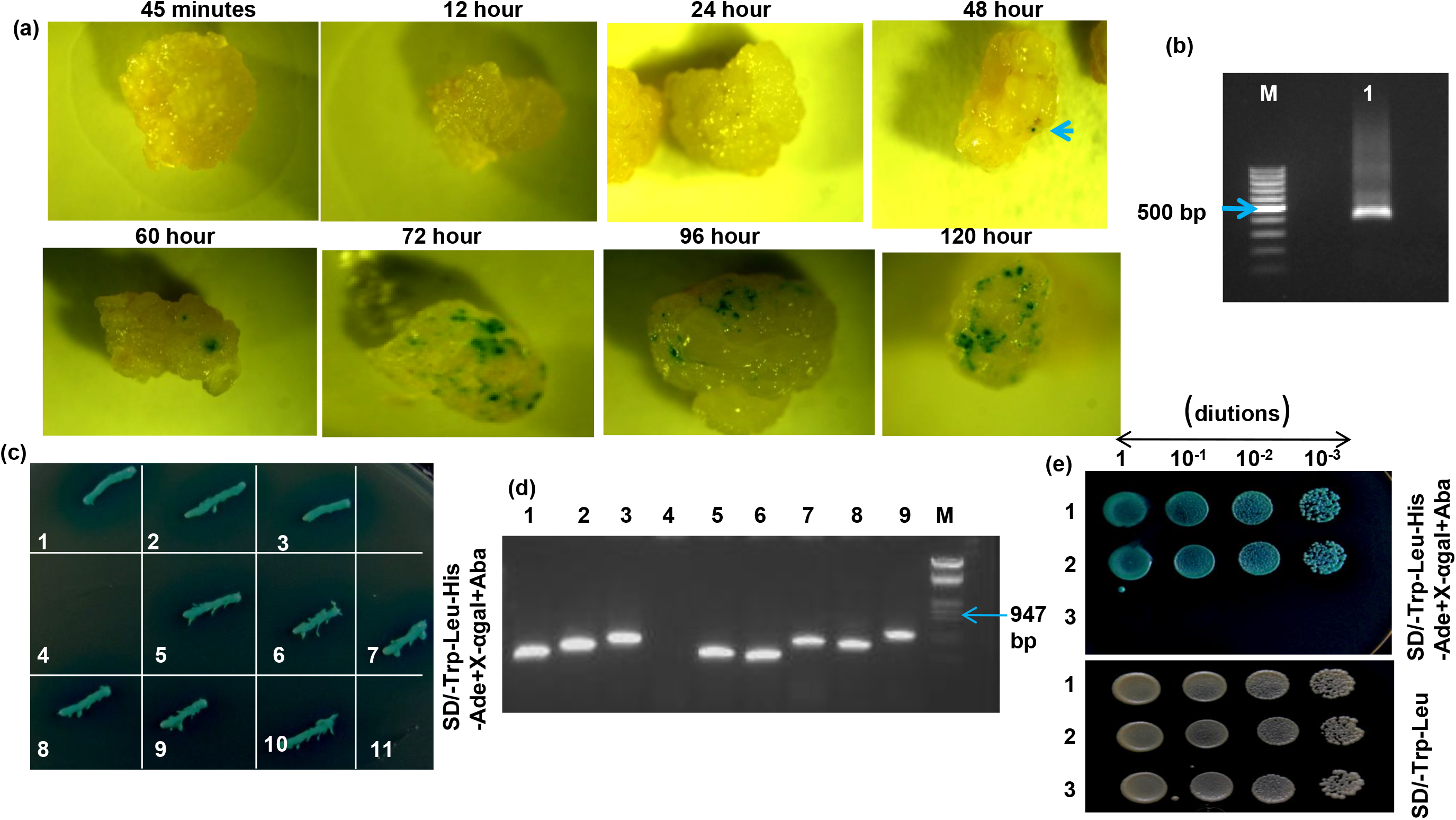
Yeast two hybrid (Y2H) assay. (a) GUS histochemical assay of rice calli inoculated with *Agrobacterium* (GUS expression cassette as T-DNA) after 45 minutes-120 hour (5^th^ day). First transient expression was observed at 48 hour (arrow). (b) 1% Agarose gel showing purified cDNA library of *Agrobacterium* infected rice calli (lane 1, lane M marker). (c) Screening of VirE2 interacting clones on SD/-Trp/-Leu/-His/-Ade/X-α-gal/Aba medium. Colony 1-9, interacting colonies, colony 10-positive control interaction (p53-BD and T antigen-AD), colony 11-negative control interaction (lamin-BD and T antigen-AD). (d) 1% Agarose gel showing PCR amplification product of interacting clones of colonies 1-9. Lane 1-9, obtained inserts, lane M-marker. (e) Specific Y2H interaction analysis between VirE2 and OsGSTU5 on SD/-Trp/-Leu and SD/-Trp/-Leu/-His/-Ade/X-α-gal/Aba mediums, row 1-positive control interaction, row 2-VirE2-BD and OsGSTU5-AD interaction, row 3-negative control interaction (VirE2-BD and T antigen-AD).

### Yeast two-hybrid (Y2H) assay

The VirE2-pGBKT7 vector construct was transformed into the *Saccharomyces cerevisiae* Y2H gold yeast strain and act as bait. The cDNA library was cloned to pGADT7-Rec vector and transformed into the *Saccharomyces cerevisiae* Y187 strain, act as prey. Yeastmaker™ Yeast Transformation System 2 kit (Clontech, Mountain View, USA) was used for the process of yeast transformation. The Y2H assay was performed following the manual provided in the kit **“**Matchmaker Gold Yeast Two-Hybrid” (Clontech, Mountain View, USA). The list of VirE2 interacting clones has been given in Fig.S1.

### Particle bombardment

2.5 µg of each purified plasmid constructs (pSAT1-nEYFP-C1-*VirE2* and pSAT1-cEYFP-C1B-*GSTU5*) were simultaneously coated on the gold particle of 1 µm size. These particles were bombarded in the onion (*Allium cepa)* peel via PDS-1000/He Biolistic Particle Delivery System model with 900 psi pressure (Sanford et al. 1993). To study the subcellular localization of *OsGSTU5*, 5 µg of the plasmid (pSAT6-EYFP-C1-*GSTU5*) construct was used for bombardment in onion peel. Peels were kept in a dark at room temperature for the flouorophore development. Peels were analyzed after 24 hour of bombardment.

### Confocal Microscopy

Fluorescence of bombarded onion peel was detected via a confocal microscope. To localize the nucleus in the peels, each bombarded peel was stained with the Hoechst 33342 (0.1-12 µg ml^-1^) for 5 minutes (for nuclear staining). Next, each peel was washed thrice with 1X phosphate buffer saline (PBS) and checked under the confocal microscopy. Confocal laser scanning microscope LSM 510 Meta was used for imaging (Zeiss, Oberkochen, Germany).

### Expression, purification, and western blot analysis of both proteins; VirE2 and GSTU5

Both pETSUMO-*VirE2* and pETSUMO-*GSTU5* constructs were transformed to BL21 (DE3) strain of the *E*.*coli* via heat-shock method. Both were analyzed for the sake of protein expression. Protein induction (20°C) was performed with 1 mM IPTG (Isopropyl ß-D-1-thiogalactopyranoside) by some modification in the protocol was given by Verma et al., 2016 (Verma et al. 2016). For immunoblotting, the PVDF membrane (Bio TraceTM 157 PVDF, 0.45 µm) was used. In order to detect a specific protein on the membrane, anti-His monoclonal antibody (Sigma, 1:1000) and anti-mouse (Sigma, 1:5000) secondary antibody were used. After tagging the primary and secondary antibody, clarity ™ Western ECL substrate (luminol enhancer solution and peroxide solution) was used to detect the blot in the ChemiDoc.

### His-pull down assay

For His-pull down assay, pETSUMO-*GSTU5* was digested with the SUMO protease (Invitrogen) to remove the His-tag from GSTU5-SUMO fusion protein. For 1 μM protein digestion, 1 unit of SUMO protease was used. Purified His tagged VirE2 and GSTU5 (SUMO digested) proteins were independently bound on Ni-NTA agarose beads for four hours (20 mM Tris, 150 mM NaCl, 1 μM β-mercaptoethanol). Beads being centrifuged, the washing and elution fraction was loaded on 12% SDS-PAGE to analyze the binding of VirE2 and GSTU5 on the bead. The VirE2 incubated beads were further incubated overnight with the GSTU5 (without SUMO tag) protein in the absence and presence of 10 mM GSH in the same buffer. The beads were washed and eluted with 250 mM imidazole. The washing and elution fractions were run on the 12% SDS-PAGE. Both the proteins were analyzed by western blot. The VirE2 protein was detected by the anti-VirE2 polyclonal antibody (G-Biosciences, 1:10000) and anti-rabbit secondary antibody (Sigma, 1:5000); GSTU5 was detected by the anti-GST primary antibody (Novex, 1:500) and anti-mouse secondary antibody (Sigma, 1:5000).

### mRNA isolation and qRT-PCR

Total RNA was isolated by QIAGEN RNeasy Plant Mini Kit (QIAGEN, USA). Besides, cDNA synthesis had been performed according to the given protocol (Kidwai et al. 2019). qRT-PCR was performed using Fast SYBR Green master Mix (ABI, USA). The relative amount of the calculated mRNA was normalized by using the rice *actin* gene (AK060893) as a control. Transcript quantification was done by the qRT-PCR using 7500 fast real-time PCR system (Applied Biosystems, USA).

### S-glutathionylation assay

To study the in vitro S-glutathionylation of VirE2 protein, 1 μM of VirE2 (A), 0.5 μM of GSTU5 (B), and 10 mM of GSH (C) added together. Four reactions were prepared **1**. A+B+C, **2**. A+B-C, **3**. A-B+C and **4**. A-B-C. All reactions were prepared in the 0.1 M phosphate buffer (pH 6.5). Reactions were kept at 30° C for 10 minutes and let run on non-reducing SDS-PAGE and transferred on the PVDF membrane. The VirE2 protein was detected by anti-VirE2 polyclonal primary antibody (G-Biosciences, 1:10 000) and anti-rabbit secondary antibody (Sigma, 1:5000). The glutathionylated VirE2 was detected by the anti-glutathione monoclonal primary antibody (My biosource, 1:1000) and anti-mouse (Sigma, 1:1000) secondary antibody (Klaus et al. 2013).

### RMSD (root-mean-square deviation) plot analysis

Protein (VirE2), ssDNA, and ligand (GSH) systems were prepared by docking them in the Hex docking tool. The docked structures were further used in molecular dynamics simulation to observe trajectory change. A 10 ns long simulation system was prepared in the space water system. The output file of trajectories was used for RMSD plots. The DNA sequence had been given in the supplementary table.

### EMSA Analysis

For 3’ end biotin labelling, 5 μM of the ssDNA (oligo) was reacted with 25 μM of Biotin 11UTP and 1.5 unit of the terminal deoxynucleotidyl transferase (TdT) which catalyzes the incorporation via non-template-directed deoxynucleotide, onto the 3’-OH end of the DNA. The reaction was kept at 37°C for 30 minutes. The reaction was stopped with 0.2 M EDTA and purified by chloroform: Isoamyl alcohol (24:1). To label the oligo with biotin, Pierce Biotin 3’ end labeling kit (PI89818), Thermo fisher scientific, was used.

The VirE2 protein was glutathionylated as described above with 10 and 20 mM GSH and purified via Ni-NTA Agarose bead. For EMSA assay, 100 ng VirE2, 100 ng glutathionylated VirE2, 100 ng GSTU5, 10 mM GSH, 20 mM GSH, were incubated with 2 ng of Biotin-experimental oligo (1X binding buffer, 2.5% glycerol, 5 mM MgCl_2_, 50 ng μl^-1^ poly (dI•dC), 0.05% NP-40) either independently or in different combinations. To achieve control reaction 20 fmol of the biotin-control oligo and 1 unit of the EBNA (Epstein-Barr Nuclear Antigen) extract were used. The reaction was kept at room temperature for 1 hour. 0.5X TBE was used as the running buffer and gel was prerunned for 1 hour at 4°C. The samples were loaded on the gel and after running 1/3^rd^ of the gel (4°C), it was transferred to the positively charged Nylon membrane (Biodyne B positively charged nylon membrane, Thermo fisher scientific). After that the blot was cross-linked via UV lamp at 254 nm for 10 minutes then processed according to the given protocol (Light Shift Chemiluminescent EMSA kit, Thermo fisher scientific). The blot was quantified by ImageJ software.

### *Agrobacterium*-mediated transformation

Empty Vector (EV), Over-Expressed *GSTU5* (OE), and Knock-Down *GSTU5* (KD) constructs were transformed to *Agrobacterium* strain EHA101 using the freeze-thaw method (Wise et al. 2006). For Agroinfiltration in transgenic rice (T_2_ generation), the method given by Andrieu (Andrieu et al. 2012) has been used with some modification.

### Rice transformation

*Agrobacterium* strain EHA101 with desired constructs in pIRS154 vector was used to transform the rice calli, nipponbare (*O*.*sativa L*. ssp *japonica*) with some modification in the given protocol (Shri et al. 2013). Three rounds of selection were done on the transformed calli (each selection for 15 days under hygromycin selection pressure). Regeneration was done according to the protocol (Chakrabarty et al. 2010). The transgenic calli were retransformed with *Agrobacterium* strain EHA101 harboring pIG121-Hm^51^ binary vector harbouring GUS expression cassette (with intron) and kanamycin resistant gene as a selection marker. The retransformed transgenic calli or leaves were selected under kanamycin resistance. The optical density of *Agrobacterium* was maintained to OD_600_ 0.8 during rice transformation.

### Screening of transgenic plants

Rice plantlets grown from transformed rice calli were transferred to the pots filled with the soilrite and hewitt media. Plantlets were first kept under the optimum physiological condition of 16 hour light (120 µmol m^-2^ sec^-1)^ and 8-hour dark photoperiod at 25±2 □C temperature and then transferred to glasshouse for growth. At the tillering stage, the genomic DNA was isolated from the flag leaf of rice plants and tested for the hygromycin gene (1026 bp) via PCR with the HptII primers. Reaction products were run on 1.2% agarose gel (Fig. S 3a, b, c, d).

### Analysis of AMT efficiency

To analyze the *Agrobacterium*-mediated transformation (AMT) efficiency in rice transgenic, the T_2_ generation lines/calli was selected according to GSTU5 transcript abundant in qRT-PCR. The T_2_ transgenic lines/calli were transformed with *Agrobacterium* strain EHA101 (having GUS expression cassette as T-DNA in a pIG121-Hm^51^ binary vector). The calli, after three days of co-cultivation, were transferred to selection medium. For selection, hygromycin (30 mg l^-1^) and kanamycin (50 mg l^-1^) drugs were used in combination. After 3 weeks of selection period the calli were analyzed for CBR, survival rate and GUS assays. To visualize the GUS expression, GUS histochemical staining (Jefferson et al. 1987) was used. To detect GUS protein total protein was isolated from the agroinfected rice calli (Rai et al. 2011). An equal amount of protein was added to the GUS assay buffer with 1 mM pNPG. The solutions were kept at 37°C for one hour, GUS protein measured spectrophotometrically at 405 nm. Callus browning rate (CBR) was calculated from total brown calli/ total transformed calli *100. The survival rate was too calculated from total survived calli/total transformed calli*100. The Agroinfiltered leaves were placed on co-cultivation media for 4 days. The transformation was calculated from total GUS expressing calli or leaves/ total transformed calli or leaves*100.

### Glutathione-S-transferase (GST) activity

To analyze the GST in transgenic plants, flag leaf (0.5 g fresh weight) from each line was grounded to powder in liquid nitrogen. The grounded leaf is placed in enzyme extraction buffer (1 ml phosphate buffer, 1% polyvinylpyrrolidone with 0.1% phenylmethylsulfonyl fluoride), centrifuged at 12000*g* for 15 minutes at 4°C. The pellet was discarded and supernatant was used to analyze the GST activity (Thounaojam et al. 2012). GST activity was performed according to method (Habig et al. 1974) by utilizing GST Assay kit (Sigma, USA, CS0410-1KT)

### Statistical analysis

Each experiment was performed thrice in randomized manner. Mean, standard error, and triplicate data were used for the statistical evaluation using GraphPad Prism5. The statistical significance of differences between samples was tested using the Duncan’s test. Significance was measured by utilizing SPSS 16.0 software at *P*≤0.05.

## Results

### Screening of VirE2 interacting protein in rice

To identify VirE2 interacting protein in rice, a cDNA library from *Agrobacterium* inoculated rice calli was generated and yeast two-hybrid (Y2H) assay was performed (Iwabuchi et al. 1993; Li and Fields 1993). We chose to construct the prey library from *Agrobacterium*-inoculated rice calli, because we were interested to find out the interacting partners that might be expressed or upregulated after perception of the *Agrobacterium* or delivery of T-DNA/virulence proteins. To select the appropriate prey for cDNA library preparation, the rice calli were inoculated with *Agrobacterium* strain EHA101 with pIG121Hm^51^ binary vector [harbouring “GUS (β-glucuronidase with intron) expression cassette as T-DNA”]. The expression of T-DNA (GUS) was assayed at different time intervals. The first expression was observed after 48 hours of inoculation (transient expression) which was subsequently increased with time upto 120 hour (Fig. 1a). A cDNA library was prepared from *Agrobacterium* inoculated rice calli (from 1, 2, and 3 day post inoculated calli) (Fig. 1b). The Y2H assay was performed between VirE2 (as bait) and cDNA library from rice calli (as prey). We identified eight clones that interact with VirE2 protein (Fig. 1c, d, Fig.S1). One of the identified clones, the tau class glutathione-S-transferase *LOC_Os09g20220* (*OsGSTU5*), was studied in depth.

The full-length cDNA sequence of *OsGSTU5* (687 bp) was obtained from the Rice genome annotation project (TIGR database). The specific interaction between full-length OsGSTU5 and VirE2 was further confirmed by Y2H assay. This distinctly showed growth on appropriate selection media and confirmed the specific interaction between VirE2 and OsGSTU5 (Fig. 1e). Interaction between murine p53 and large T-antigen cloned in pGBKT7 (BD) and pGADT7 (AD) vectors, respectively, was taken as the positive control mating. The interaction between murine p53-BD and lamin-AD was taken as the negative control mating during Y2H screening (Fig. 1c, d). During specific interaction, the positive control was the same but negative control interaction was performed in between VirE2-BD and T antigen-AD (Fig. 1e).

### VirE2-OsGSTU5 interaction occurs in plant cytoplasm

To demonstrate the VirE2-OsGSTU5 interaction in planta, we performed a bimolecular fluorescence complementation assay (BiFC) and analyzed the reconstituted fluorescence signal (YFP). We co-expressed the nEYFP-VirE2 and cEYFP-OsGSTU5 constructs together on onion peel via particle bombardment. The reconstituted YFP signal was observed throughout the cytoplasm of onion cell (Fig. 2a, panel 2). During this analysis, the EYFP construct with a complete YFP cDNA under the control of 35S promoter, was used as positive control. We observed YFP signals in the cytoplasm as well as in the nucleus (Fig. 2a, panel 1) in positive control. Another tau class GST (*Os10g38600*; OsGSTU30) from rice was utilized as a negative control during BiFC analysis (Kudla and Bock 2016). The nEYFP-VirE2 and cEYFP-OsGSTU30 constructs were co-bombarded on onion peel and analyzed for the reconstitution of the YFP signal. We did not observe the YFP signals in onion peel (Fig. 2a, panel 3). This observation further confirmed the specific interaction between OsGSTU5 and VirE2 in cytoplasm of the plant cell.

**Fig 2.**
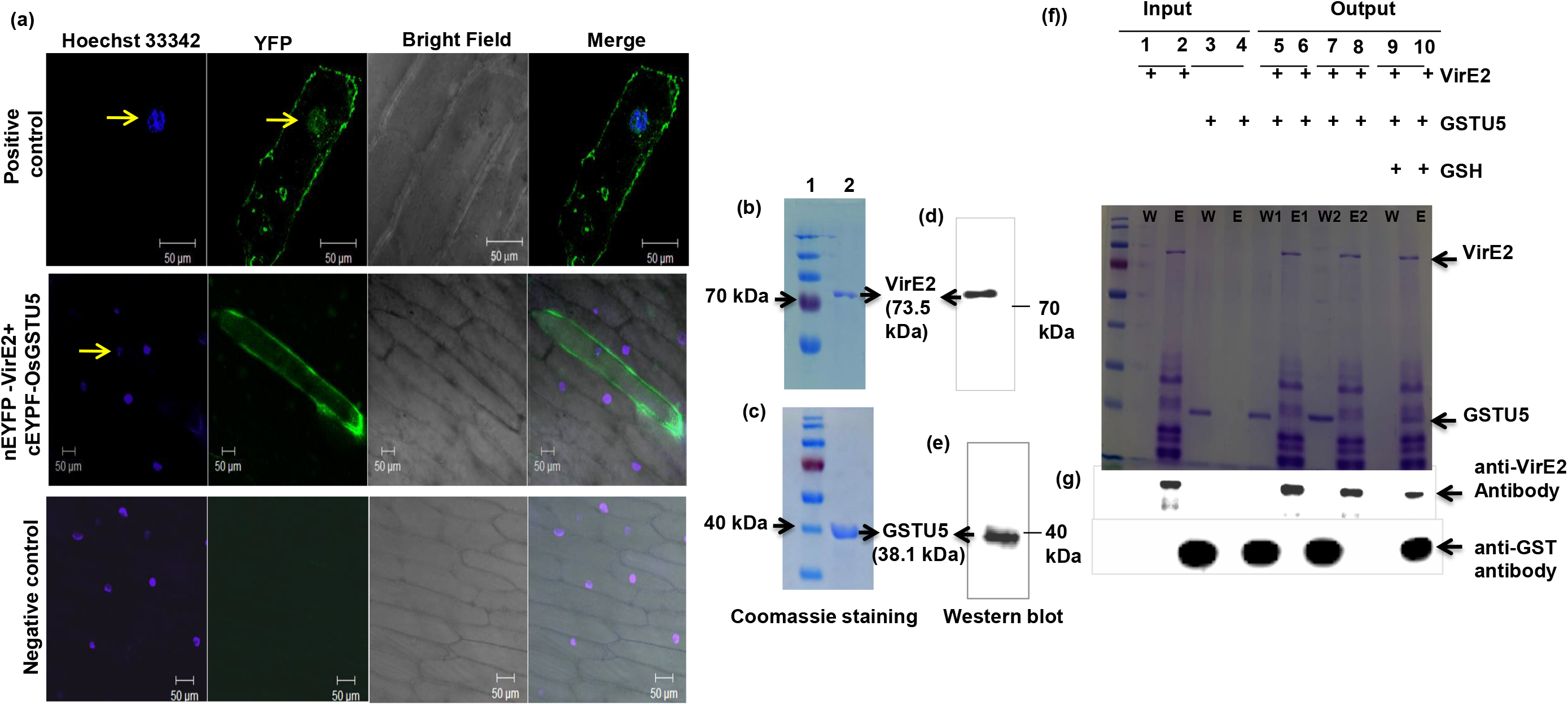
Bimolecular Fluorescence complementation (BMFC) assay and His-pull down assay. (a) Confocal image showing cytoplasmic subcellular localization of nEYFP-VirE2 and cEYFP-GSTU5 (panel 2). Positive control (EYFP) shows signal in cytoplasm and nucleus (panel 1). nEYFP-VirE2 and cEYFP-GSTU30 fusion proteins were used as negative control (panel 3). In all cases, Nuclei was stained by Hoechst 33342 (arrow). Bar 50 μM, Magnification 20X. Commassie stained SDS-PAGE of (b) purified VirE2 protein (d) purified OsGSTU5 protein. Western blot detection of (c) VirE2, (e) OsGSTU5 proteins with anti-His antibody. (f) SDS-PAGE analysis of pull down proteins, W-washing fraction, E-elution fraction, E1, E2-elution fraction 1, 2. W1,W2-Washing fraction 1 and 2. (g) Western blot detection of pull down proteins.

### In vitro VirE2-OsGSTU5 interaction requires glutathione

VirE2 and OsGSTU5 were both expressed as His-tagged fusion proteins in *E. coli* and purified on Ni-NTA agarose beads. Both purified proteins were analyzed on the SDS-PAGE via coomassie staining, and also detected via the respective antibodies (Fig. 2b, c, d and e). Later, the SUMO tag was removed from the GSTU5-SUMO fusion protein and analyzed on the SDS-PAGE via coomassie staining (Fig. S2a).

We analyzed the *in vitro* VirE2-GSTU5 interaction by performing a pull-down assay. To confirm the indirect binding of GSTU5 on Ni-NTA agarose bead, the GSTU5 purified protein (without SUMO or His tag) was incubated with the bead. We observed all the GSTU5 in washing fraction (W) and not in the elution fraction (E) either in SDS-PAGE or western blot (Fig. 2f, g lane 3, 4). This observation confirms the absence of indirect binding of GSTU5 with the bead. Further, VirE2 was also incubated with bead and the VirE2 fraction was observed in E fraction (Fig. 2f, g lane 1, 2), which confirms the binding of VirE2 protein with the bead. For pull-down analysis, VirE2 and GSTU5 proteins were incubated on bead in presence and absence of glutathione (GSH). We have analyzed two pull-down fractions of VirE2-GSTU5 interaction conducted without GSH. We observed the GSTU5 only in W fraction and VirE2 in E fraction (Fig. 2f, g lane 5, 6, 7, 8). This clearly confirms that GSTU5 is not able to interact with VirE2 protein in the absence of GSH. Further, VirE2 and GSTU5 were bound together on the bead in presence of 10 mM GSH. Here we found that both VirE2 and GSTU5 were eluted together in E fraction and no GSTU5 was observed in W fraction (Fig. 2f, g lane 9, 10). The observation convincingly confirms the in vitro interaction between VirE2 and GSTU5 in presence of GSH.

### GSTU5 is localized in the plant cytoplasm and has a higher transcript level in *Agrobacterium* infected rice calli

After confirmation of the interaction between GSTU5 and VirE2 in the cytoplasm of the plant cell (Fig. 2a, panel 2), we observed the subcellular localization of the GSTU5 by using a GSTU5-YFP fusion protein. We have used a particle bombardment tool to transiently express the GSTU5-YFP fusion protein on onion peel. After fluorophore development, we analyzed the YFP signals in onion peel by confocal laser microscopy. Though we observed the YFP signal throughout the cytoplasm, it was not found in the nucleus (Fig. 3, panel 2). This observation reaffirms that GSTU5 localized in the plant cytoplasm. Here, the same positive control was used as in BiFC analysis (Fig. 3, panel 1). For negative control, cEYFP-GSTU5 construct was used (Fig. 3, panel 3).

**Fig 3.**
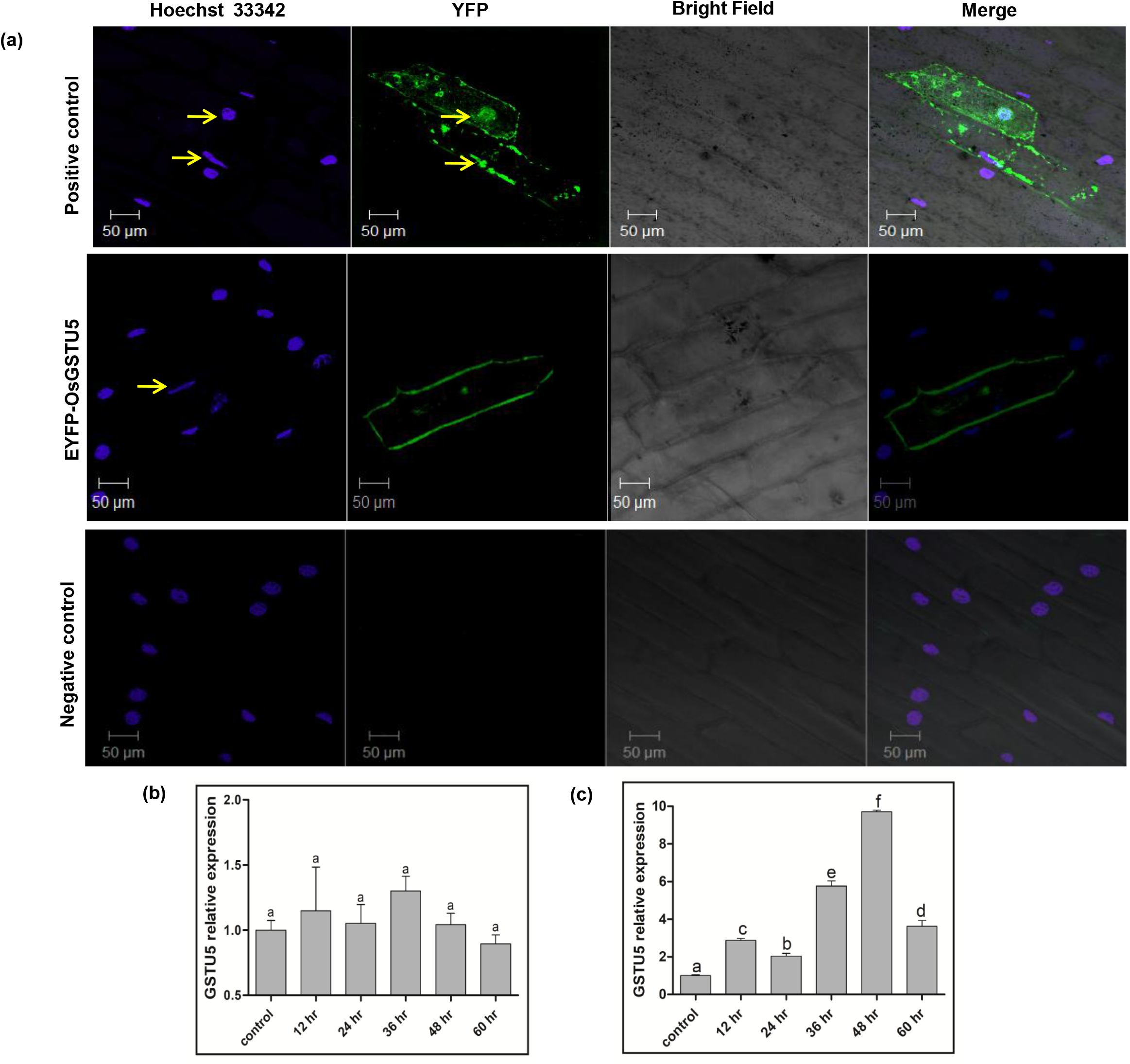
OsGSTU5 localization and relative expression. (a) Confocal image showing EYFP-GSTU5 subcellular localization in cytoplasm (panel 2), EYFP was used for positive control which show signals in cytoplasm and nucleus (panel 1), cEYFP-GSTU5 was used as negative control (panel 3). In all cases, Nuclei was stained by Hoechst 33342 (arrow). Bar 50 μM, Magnification 20X. Relative expression of *OsGSTU5* in (b) mock and (c) *Agrobacterium* inoculated rice calli. The rice calli placed on same co-cultivation medium without *Agrobacterium* infection was considered as mock.

The expression profile of *GSTU5* transcripts was also analyzed in rice calli. We inoculated the rice calli with and without *Agrobacterium*. The rice calli without *Agrobacterium* inoculation on the same co-cultivation media was considered as mock. The modulation in the *GSTU5* transcript at different time intervals was analyzed in these rice calli. No significant difference in the GSTU5 transcript was traced in mock rice calli (Fig. 3b) whereas an upregulated *GSTU5* transcript was observed in the rice calli with *Agrobacterium* inoculation. We observed the highest *GSTU5* transcripts in 48 hr post-inoculated calli (Fig. 3c).

### In vitro analysis confirms the glutathionylation of VirE2 protein by GSTU5

S-glutathionylation of protein is an important post-translational modification that introduces a negative charge on protein and may alter the protein function (Dalle-Donne et al. 2009). An in vitro S-glutathionylation assay was performed to analyze the glutathionylation of VirE2 protein. The anti-VirE2 antibody was used to detect VirE2 protein and the anti-glutathione antibody was used to detect glutathionylated VirE2 (gVirE2) protein. VirE2 was detected in all lanes, while gVirE2 was detected only in one lane in which GSTU5 and GSH were present (Fig. 4a, blot 1 and 2). Thus the in vitro S-glutathionylation assay confirms that GSTU5 glutathionylates the VirE2 protein.

**Fig 4.**
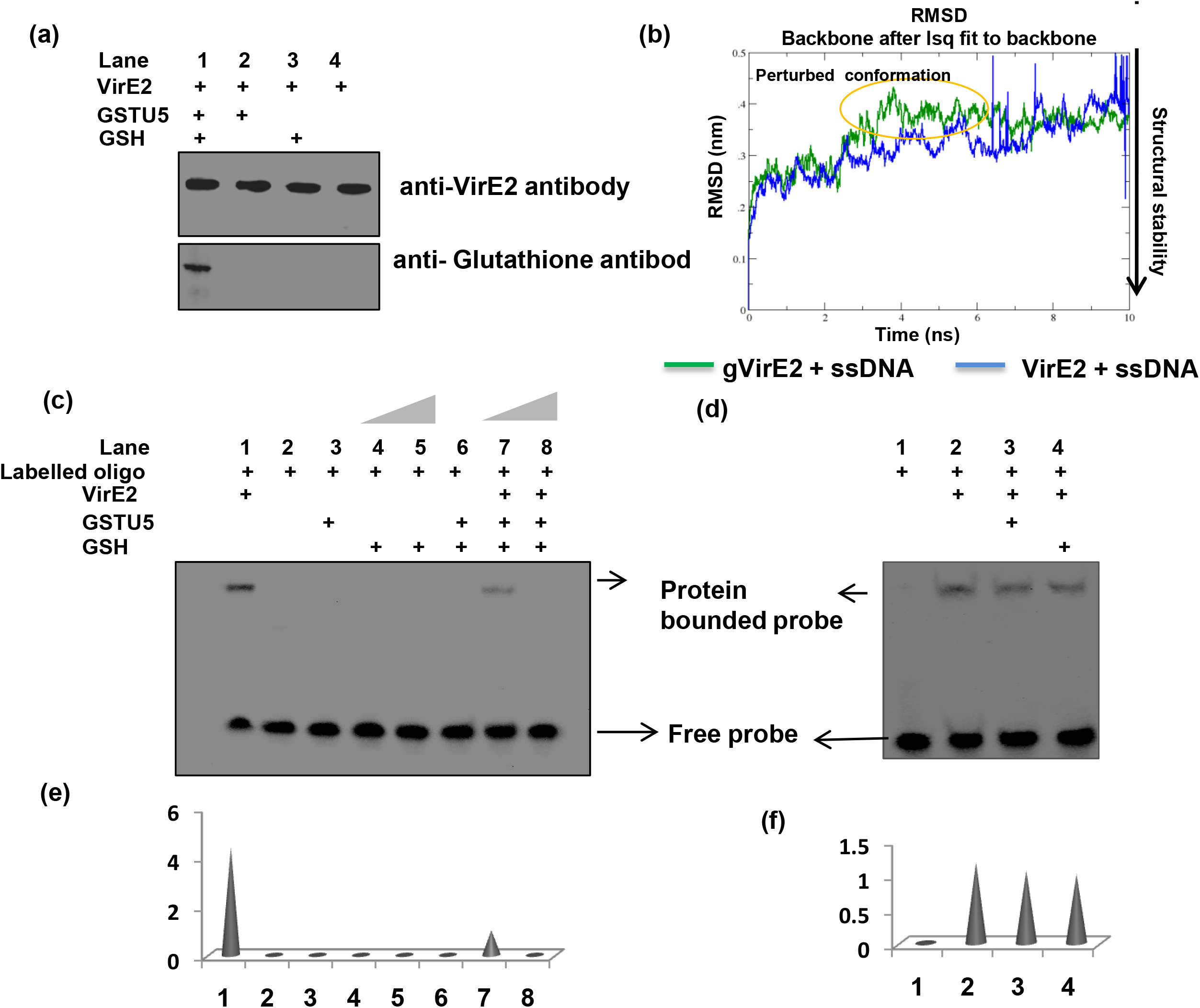
*In vitro* analysis of S-glutathionylation and ssDNA binding activity of VirE2 protein: (a) S-glutathionylation assay-VirE2 protein was incubated with GSTU5 and GSH in different combinations and VirE2 was detected with anti-VirE2 antibody (blot 1, lane 1-4). Glutathionylated VirE2 (gVirE2) was detected by anti-glutathione antibody only in one lane (blot 2, lane 1). (b) RMSD plot analysis of “VirE2+ssDNA” Versus “gVirE2+ssDNA” complex. (c) EMSA assay-Analysis of SSB of VirE2, GSH, GSTU5 either independently or in combination with Biotin-experimental oligo (ssDNA, 39 nucleotide long), probe was found to be bounded only with VirE2 protein (Shift in lane 1). The ssDNA binding property of VirE2 protein was decreased after glutathionylation with 10 mM GSH (shift in lane 7). No shift was observed when VirE2 was glutathionylated with 20 mM GSH (lane 8). A wedge over lane 4, 5, 7, 8 denotes the 10 and 20 mM GSH concentration, respectively (d) Analysis of SSB activity of VirE2, VirE2+GSTU5 and VirE2+GSH. (e) Quantification of band intensity of protein bound probe of image (c). (f) Quantification of band intensity of protein bounded probe of image (d).

### “gVirE2+ssDNA” complex shows structural and conformational instability

To analyze the structural and conformational stability of the protein-DNA complex, we performed the MD (molecular dynamics) simulation of “VirE2+ssDNA” and “gVirE2+ssDNA” complexes. The structural stability of the constructed model was analyzed by the root-mean-square deviation (RMSD) plot. The RMSD plot was analyzed via MD simulation trajectory base analysis for 10 nanoseconds (ns) (Fig. 4b). The first 2 ns were considered as an equilibrium system for both the complexes. We observed that after 3 ns the “gVirE2+ssDNA” complex showed a change in structural trajectory in comparison to standard system (VirE2+ssDNA). It could be due to the alteration in structural geometry of gVirE2 protein which tends to increase the RMSD (Fig. 4b). On the other hand, the standard system also showed some high peaks in the RMSD graph. These peaks indicated atom-wise structural fluctuations because of structural stabilization. It was noted that, despite those fluctuations, the aforesaid complex maintained a persistent converged state. This conformational dynamics-based study indicates that the “VirE2+ssDNA” complex is more stable in comparison to the “gVirE2+ssDNA” complex.

### gVirE2 attenuates its ssDNA binding property

VirE2 has been reported to serve as SSB protein which binds with T-DNA (ssDNA) and assists T-DNA cytoplasmic trafficking and perhaps the nuclear import (Citovsky et al. 1989; Rossi et al. 1996; Yang et al. 2017; Yusibov et al. 1994). Through RMSD plot analysis we analyzed the structural instability of “gVirE2+ssDNA” complex. Next, we tested the ssDNA binding property of VirE2 and gVirE2 by electrophoretic mobility shift assay (EMSA). A biotin-labelled 39 nucleotides long single-stranded DNA (experimental oligo) was used to analyze the gel shift. After confirmation of the labelling and control reactions (Fig. S2b), we checked whether the other components of glutathionylation, like GSTU5 or GSH, had the same SSB property or not. For this, we analyzed the ssDNA binding activity of GSTU5, 10 mM GSH, 20 mM GSH, GSTU5+GSH with experimental oligo. We did not identify any shift in the experimental oligo. This finding confirms that neither GSTU5 nor GSH binds with ssDNA (Fig. 4c, lane 3, 4, 5, 6). Further, both VirE2 and gVirE2 were assessed for their SSB property. An intense shift was observed when VirE2 was incubated with a probe (Fig. 4c, lane 1). This shift confirms that VirE2 protein binds with ssDNA. The intensity of the shift was found successively decreasing when the probe was incubated with gVirE2 (Fig. 4c, lane 7, 8). We also tested the T-DNA binding property of VirE2, VirE2+OsGSTU5, VirE2+GSH through gel shift. We did not observe any significant difference in the T-DNA binding property of VirE2 protein (Fig. 4d, lane 2, 3, 4). To check the intensity of shift we used the imageJ software which showed a significant difference in VirE2-DNA and gVirE2-DNA complexes (Fig. 4e). Significantly, no notable difference was observed in the intensity of gel shift when the probe was incubated either with GSTU5 or GSH in combination to VirE2 protein (Fig. 4f). The observation suggested that gVirE2 loses its SSB property in comparison to VirE2 protein. Previous reports suggested an increased intracellular GSH concentration during biotic stress in plant (Noctor et al. 2002). It can be correlated that during *Agrobacterium* infection the plant will maintain high GSH content to perform the glutathionylation process.

### Generation of *OsGSTU5* transgenic plant with modulated GST activity under control condition

To characterize functionally the role of GSTU5 in planta, we cloned the Over-Expressing (OE) and Knock-Down (KD) *OsGSTU5* constructs under the control of Ubiquitin maize promoter in pIRS154 vector. All the constructs were transformed in *Agrobacterium* strain EHA101. Further, we generated the pIRS154 or Empty Vector (EV), Over-Expressing (OE) and Knock-Down (KD) *OsGSTU5* transgenic rice plants by transforming the rice calli with respective *Agrobacterium* constructs. All the constructs also contained hygromycin phosphotransferase (*HPTII*) marker gene along with the desired gene. For initial screening, the genomic DNA from all transgenic rice plants was tested for the presence of *HPTII* marker gene through PCR (Fig. S3 a, b, c, d). At first we analyzed the three independent lines of both, i.e., OE and KD for *OsGSTU5* transcripts via qRT-PCR. The OE lines had 3.3, 2.2 and 2.0 fold *GSTU5* transcripts in OE-1 and OE-9 and OE-10 lines, respectively compared to EV. The KD lines had 0.10, 0.40 and 0.44 fold *GSTU5* transcripts respectively in KD-20, KD-22 and KD-25 compared to EV (Fig. 5a). The data suggested the upregulation of *GSTU5* transcripts in overexpression lines while downregulation in KD lines.

**Fig 5.**
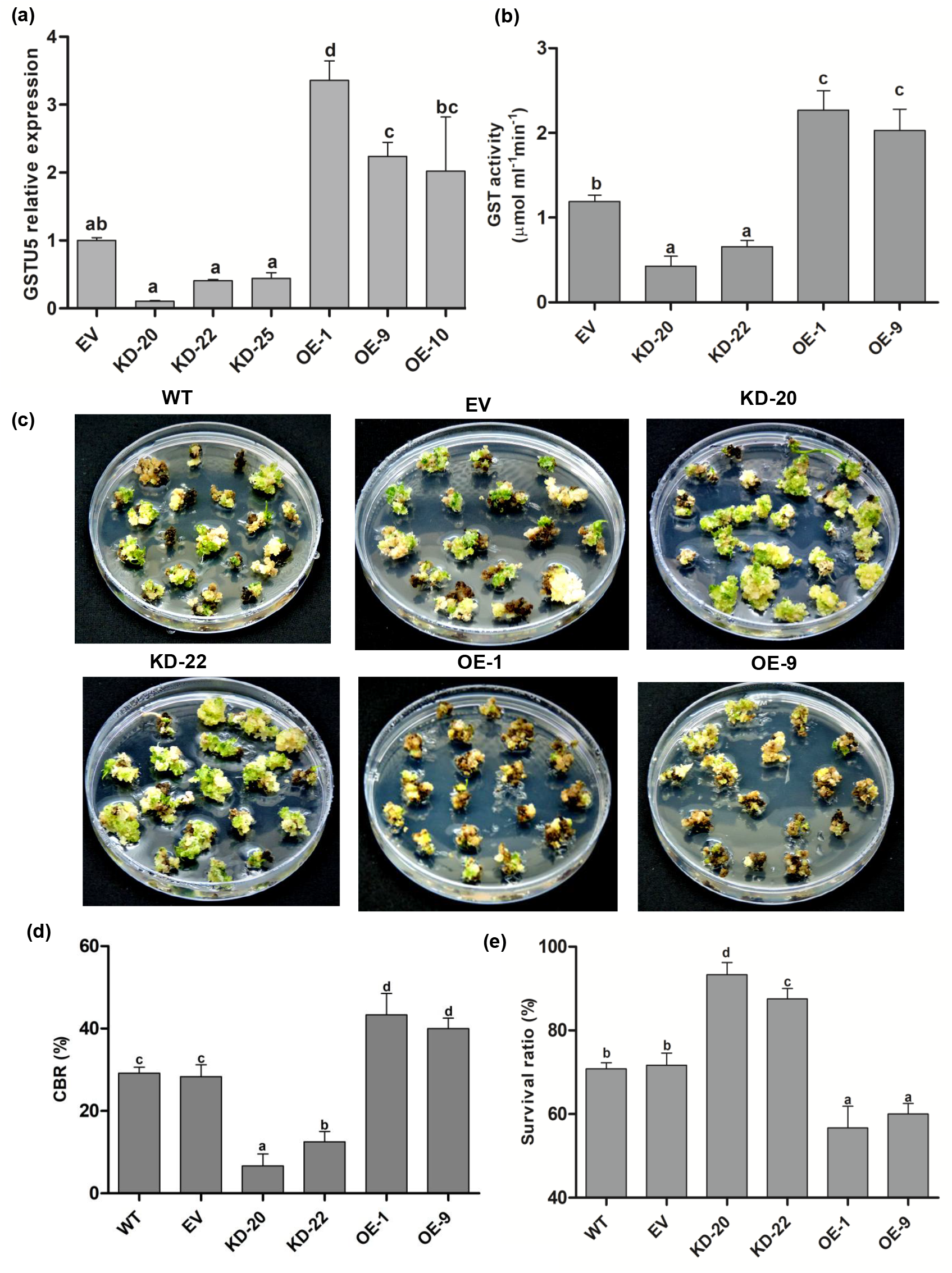
Confirmation of transgenic rice plants and AMT efficiency in rice calli. (a) *GSTU5* relative expression and (b) GST activity in transgenic rice plants at T_2_ generation. (c) Rice calli (WT and transgenic) on selection medium, (d) Callus browning rate (CBR) and (e) Callus survival rate after 3 weeks post *Agrobacterium* infection. Values are mean ±SE (n=3): Different letters shows the significant difference while same letter shows no significant difference, (Duncan, *P*≤0.05).

To further confirm the transgenic lines, two best expressing lines OE-1, OE-9, and KD-20, KD-22 were analyzed for GST activity under control condition. We found approximately 1.9 and 1.7 fold higher GST activity in OE-1 and OE-9 lines, respectively and an approximately 50% decrease in GST activity in KD lines in comparison to EV (Fig. 5b). The higher and lower GST activities in OE and KD lines can respectively be correlated with overexpression and down-regulation of *OsGSTU5*, in these lines.

The transgenic lines, as well as wild type (WT) plant, were tested for the seed viability and plant morphology. Here we did not observe any significant difference among all lines including WT (Fig. S4, a, b, c, d). It confirms that *OsGSTU5* does not interfere with the essential physiological process in the rice plant.

### Knock-down lines were prone to *Agrobacterium*-mediated transformation/infection

We further tested the ability of OE and KD *OsGSTU5* rice calli for their susceptibility during *Agrobacterium* infection. We first tested the probable effect of the transgenic background on the subsequent *Agrobacterium*-mediated transformation/infection. We used EHA101 *Agrobacterium* strain with pIG121-Hm^51^ binary vector for subsequent transformation of transgenic rice calli. The pIG-121Hm^51^ binary vector possessed GUS expression cassette (with intron), neomycin phosphotransferase II) *NPTII* and (hygromycin phosphotransferase II) *HPTII* genes that provide tolerance against kanamycin and hygromycin drugs, respectively (Shri et al. 2014, Tiwari et al. 2020). We generated the calli from WT, EV, KD and OE lines and transformed with EHA101 (pIG-121Hm^51^ as binary vector). After transformation, the calli were shifted to selection medium with kanamycin drug as selection marker. We did not identify any significant difference between WT and EV line for their *Agrobacterium* infectivity. It clearly indicates that the procedure used for transgenic generation did not make them resistant to subsequent *Agrobacterium* infection (Fig. 5c). The callus browning ratio (CBR) is a good indicator of defense response of calli during *Agrobacterium*-mediated gene transfer (Zhang et al, 2020), therefore we analyzed the CBR in all lines. The KD calli with ∼10% callus browning ratio (CBR) and ∼90% survival in comparison to WT/EV showed they were highly susceptible for *Agrobacterium* infection. At the same time, the OE lines with ∼40% CBR and ∼60% survival ratio in comparison to WT/EV plant confirmed their resistance during *Agrobacterium* infection (Fig. 5d, e).

### Contrasting T-DNA expression in OE and KD lines

The inefficiency of VirE2 protein to bind with T-DNA will ultimately hamper T-DNA journey, integration and thus expression in the transgenic plant. Keeping this in mind, we retransformed the transgenic leaves and calli with *Agrobacterium* strain EHA101 (pIG121-Hm^51^ as binary vector). The retransformed transgenic calli selected on kanamycin drug for 3 weeks and analyzed for stable T-DNA/GUS integration (Janssen et al.1990). For transient T-DNA expression analysis, the GSTU5 transgenic leaves were retransformed with same *Agrobacterium* strain. The GUS expressing leaves were analyzed after 4 days of co-cultivation (Janssen et al.1990).

We found a drastic change in GUS (T-DNA) expression in the transgenic rice calli. The KD lines had a higher number of GUS expressing calli in comparison to EV. In contrast, the OE lines had a comparatively lower number of GUS expressing calli (Fig. 6a). The total GUS transcript was approximately four-fold upregulated in KD calli while it was found five-fold downregulated in OE calli (Fig. 6b). The GUS activity in these lines was also calculated. We observed ∼50% decreased GUS activity in OE lines than that of KD lines (Fig. 6c). T-DNA transfer efficiency was also calculated which was ∼80% in KD lines followed by ∼60% in EV. The least transformation with ∼40% efficiency was observed in OE lines (Fig. 6d).

**Fig 6.**
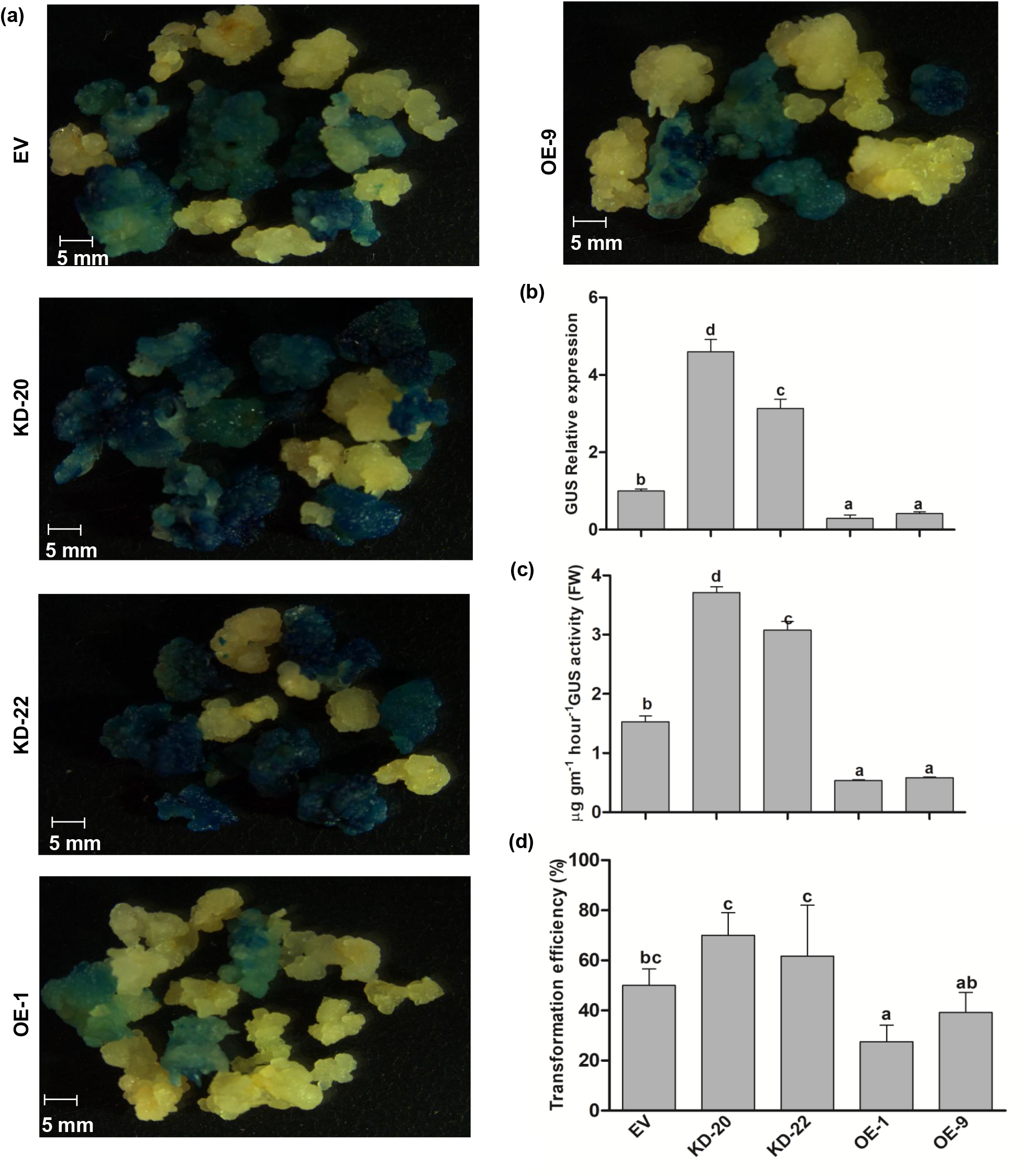
T-DNA (GUS) expression in rice calli. (a) GUS histochemical assay, (b) GUS relative expression, (c) GUS activity, and (d) Transformation efficiency in transgenic rice calli after 3 weeks post *Agrobacterium* (GUS expression cassette as T-DNA) infection. Values are mean ±SE (n=3): Different letters shows the significant difference while same letter shows no significant difference, (Duncan, *P*≤0.05).

Further, we agroinfiltered the greenhouse-grown transgenic rice leaves with the same *Agrobacterium* strain. For the total analysis, ten leaves from each line were analyzed. We observed higher number of GUS expressing leaves in agroinfiltered KD lines. Besides, these lines possessed a higher GUS intensity with 2.5 fold upregulated GUS activity and ∼80% GUS expressing leaves. The least efficiency with a ∼40% GUS expressing leaves was observed in OE lines (Fig. 7a, b, c). In brief, we observed the highest T-DNA expression in KD lines followed by EV, and the least expression was observed in OE lines.

**Fig 7.**
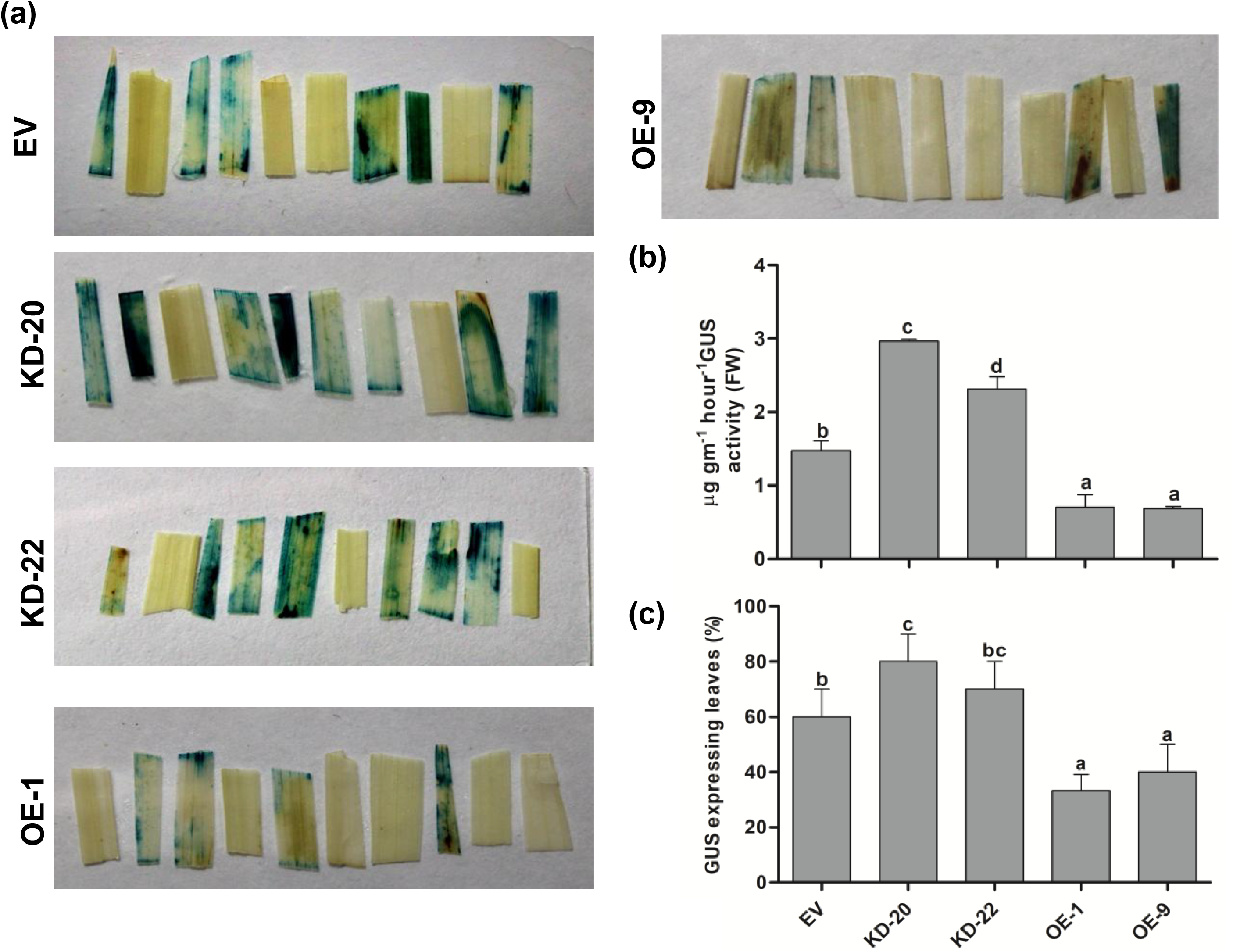
T-DNA (GUS) expression in rice leaves. (a) GUS histochemical assay, (b) GUS activity, (c) GUS expression efficiency of transgenic rice leaves after 4 days of *Agrobacterium* (GUS expression cassette as T-DNA) infection (Agroinfiltration). Values are mean ±SE (n=3): Different letters shows the significant difference while same letter shows no significant difference, (Duncan, *P*≤0.05).

## Discussion

During AMT, the journey of T-complex through cytoplasm of the plant cell is a key regulatory event that is governed by both virulence and host proteins. Yang et al. 2017 demonstrated that VirE2 utilizes host machinery for cytoplasmic trafficking (Yang et al. 2017). Also, in vitro studies suggested that VirE2 cooperatively binds with T-DNA throughout its length (Christie et al. 1988; Citovsky et al. 1989). Therefore, it can be hypothesized that VirE2 protein assists the cytoplasmic trafficking of T-DNA.

On note, no host protein from monocot plant has yet been reported to interact with VirE2 protein during AMT. Here we identified eight host proteins of rice that interact with VirE2 protein, including a tau class GST, OsGSTU5 (Fig.1, Fig. S1.). GST serves as a potent detoxifying enzyme (Hayes and Pulford 1995) and its significant role has been well-documented in the detoxification of microbial toxins. Previous reports also suggested that some of the *Arabidopsi*s mutants, resistant to AMT viz. rat (resistant to *Agrobacterium* transformation), have higher transcripts of early and late defense genes in contrast to wild-type. The wild-type only has early defense genes, and no late defense genes were observed (Zhu et al. 2003). Veena et al. (2003) identified various types of glutathione-S-transferases and other defense-related genes differentially expressed during the *Agrobacterium*-plant interaction (Veena et al. 2003). It was also reported that *Agrobacterium*-mediated transformation in monocot cells was directly associated with the defense response of plants (Zhang et al. 2013). The above premises prompted us to select *OsGSTU5* (*Os09g20220*) for detail analyses in this study.

The localization study confirms the VirE2-OsGSTU5 interaction in cytoplasm of the plant cell (Fig. 2a, panel 2). Also, we found the upregulated *GSTU5* expression in rice calli during *Agrobacterium* infection (Fig. 3c). These observations confirm the probable role of GSTU5 in host cytoplasm during *Agrobacterium* infection. Further, the in vitro analysis confirmed that GSH is required during VirE2-OsGSTU5 interaction (Fig. 2f, g), suggesting the glutathionylation of probable target (VirE2) protein. We confirmed this by analyzing the glutathionylated VirE2 (gVirE2) protein with anti-glutathione antibody in western blot (Fig. 4a). Previous in vitro assays confirmed that VirE2 acts as SSB protein of ssT-DNA (Citovsky et al. 1989; Rossi et al. 1996; Yusibov et al. 1994). Thus we tested the structural stability of “gVirE2+ssDNA” versus “VirE2+ssDNA” complex via the RMSD plot. The “gVirE2+ssDNA” complex was found to have perturbed protein conformation in comparison to “VirE2+ssDNA” complex (Fig. 4b). Also, SSB activity of VirE2 and the gVirE2 proteins was compared via mobility shift assay. By comparing the intensities of the shift we found that gVirE2 subsequently decreases its SSB activity while increasing the GSH concentration (Fig. 4c).

To characterize the functional role of GSTU5 during *Agrobacterium* infection in rice calli, we generated transgenic rice plants with modulated *GSTU5* transcripts. For this purpose we selected two distinct opposites - The two lines with upregulated *OsGSTU5* transcripts along with higher GST activity (OE) and the two lines with downregulated *OsGSTU5* transcript along with lower GST activity (KD) (Fig. 5a, b). We comparatively analyzed the transgenic rice calli and leaves for AMT efficiency via different experiments like CBR, survival ratio, GUS activities, transformation efficiency, and Agroinfiltration efficiency. We observed that KD lines had better transformation efficiency in comparison to EV, whereas the OE lines were least prone to *Agrobacterium* infection/transformation (Fig. 6, 7).

In a nutshell, from in vitro studies, the glutathionylated VirE2 was found to attenuate its SSB or T-DNA binding activity and in planta studies confirm the resistance and susceptibility of GSTU5 OE and KD lines, respectively during *Agrobacterium* infection. Here we propose that GSTU5 is a defense-related gene that gets activated during *Agrobacterium* infection in rice. GSTU5 by interacting and glutathionylating the VirE2 protein hampers its SSB activity. The contrasting AMT efficiency in OE and KD transgenic rice calli is possibly caused by the attenuation of VirE2 activity in the plant cytoplasm.

## Conclusion

In this study we identified that GSTU5 from rice interacts with VirE2 protein of *Agrobacterium*. GSTU5 mediated glutathionylation of VirE2 protein hampers its SSB activity. In planta studies confirmed that high GSTU5 expressing lines was less prone for *Agrobacterium*-mediated gene transfer while knocked down lines were more prone for the same. Overall, GSTU5 might act as negative regulator for VirE2 protein during AMT in rice and might utilize to improve the transformation efficiency in rice.

## Acknowledgement

The authors would like to acknowledge the Director CSIR-NBRI for the facilities to carry out this work. M.T. is indebted to Banaras Hindu University for the registration (Sept.2014/271) and UGC for providing fellowship. The authors are thankful to Dr. Emmanuel Guiderdoni, CIRAD, France, for pIRS vector and addgene for pNW55 vector. We would like to acknowledge Yogeshwar Vikram Dhar, CSIR-NBRI for *in silico* analysis.

## Author Contribution

D.C designed, supervised the experiments and reviewed the manuscript. M.T performed the experiments, executed data analysis and wrote the manuscript. A.K.M. supervised during the work. N.G., Y.I. and M.K. helped during the experimental work.

## Supplementary Information

Supplementary information includes four Figures and one table.

