## Supplementary table for "A tau class Glutathione-S-Transferase (OsGSTU5) acts as a negative regulator of VirE2 interaction with T-DNA during *Agrobacterium* infection in rice"

**Table S1- List of primers**

| | |
|---|---|
| **Primers used for Y2H assay (Sequence (5'-3')** | |
| GSTU5 (pGAD) F EcoRI | ATAGAATTCATGGCGGACGAGGT |
| GSTU5(pGAD)R BamHI | ATAGGATCCCTACTTGGCGCCAAACTTG |
| VirE2 pGBK F infusion | AGGAGGACCTGCATAATGGATCCGTCTAGCAATG |
| VirE2 pGBK R infusion | TAGTTATGCGGCCGCTCAAAAGCTGTTGACGCTTT |
| **Primers used for BifC and localization assay (Sequence (5'-3')** | |
| VirE2 ( BiFC) F XHOI | CTCGAGATGGATCCGTCTAGCA |
| VirE2 ( BiFC) R KPNI | GGTACCTCAAAAGCTGTTGACGCT |
| GSTU5 ( BiFC) F EcoRI | ATAGAATTCATGGCGGACGAGGT |
| GSTU5 ( BiFC) R BamHI | ATAGGATCCCTACTTGGCGCCAAACTTG |
| GSTU30 (BiFC) F Hind III | AAGCTT ATGGCAGGAGGAGGAG |
| GSTU30 (BiFC) R Kpn I | GGTACCTCAGTTGTTTGCGGCGGT |
| **Primers used for protein expression (pETSUMO) (Sequence (5'-3')** | |
| GSTU5 F (pET) | ATGGCGGACGAGGTGGTGCT |
| GSTU5 R (pET) | CTACTTGGCGCCAAACTTG |
| VirE2 F (pET) | CGAGTCGACATGGATCCGTCTAGC |
| VirE2 R (pET) | GAGGTCCTCGAGTCAAAAGCTGTTG |
| **Primers used for rice transgenic (Sequence (5'-3')** | |
| ImiR- s | agTTAGATATAGATCATACGCGGcaggagattcagtttga |
| IImiR-a | tgCCGCGTATGATCTATATCTAActgctgctgctacagcc |
| IIImiR*s | ctCCGCGAATGTTCTATATCTAAttcctgctgctaggctg |
| IVmiR*a | aaTTAGATATAGAACATTCGCGGagagaggcaaaagtgaa |
| GSTU5 (IRS) F Bam HI | GGATCCATGGCGGACGAGGTGGT |
| GSTU5 (IRS ) R Kpn I | GGTACCCTACTTGGCGCCAAACTTG |
| G-4368 | CTG CAA GGC GAT TAA GTT GGG TAA C |
| G-4369 | GCG GAT AAC AAT TTC ACA CAG GAA ACA G |
| G11491 | TCGGATCCCAGCAGCAGCCACAGCAAA |
| G11494 | TACCGTAGTCGTAGTCGTCGCCATGGCT |
| **Real-time primers (Sequence (5'-3')** | |
| GSTU5 F | CATCCTCGAGTACATCGACGAGACATG |
| GSTU5 R | CTGAGCTTCCAGAGGCGTGTCT |
| GUS F | AGCGCACTTACAGGCGATTA |
| GUS R | GGCACAGCACATCAAAGAGA |
| Rice Actin F | GAGTATGATGAGTCGGGTCCAG |
| Rice Actin R | ACACCAACAATCCCAAACAGAG |
| **DNA sequence used for EMSA analysis** | |
| 39 bases ssDNA | GCGGTGGAAGTAGCAACAGCGGTTCTAGCGGCAGCGGCG |
| ssDNA (used in RMSD) | CGCGAAGCATTCGCG |
| **Primers used for rice transgenic analysis (Sequence (5'-3')** | |
| HPTII F | ATGAAAAAGCCTGAACTCACCG |
| HPTII R | CTATTCCTTTGCCCTCGGACG |
| UBI F (promoter) | GTTGGGCGGTCGTTCATTCGTTCT |

**Table S1- List of primers**

| | |
|---|---|
| NosT (terminator) | GGACTCTAATCATAAAAACCC |
| **Other primers used for cloning and sequencing** | |
| F1 GSTU5 | CTTCTCCGACGAGCAGCTAG |
| F2 GSTU5 | GCAATGGCGGACGAGGTGGT |
| R1 GSTU5 | CGATTGTTAACCAAACCAAATCTGG |
| R2 GSTU5 | CACTCTACTTGGCGCCAAACTTG |
| M13F | GTAAAACGACGGCCAGT |
| M13R | CAGGAAACAGCTATGAC |
| pSAT Camv promoter | GGCTCCTACAAATGCCATCA |
| pSAT Camv terminator | CTGGGAACTACTCACACATTATTC |
| T7 promoter | TAATACGACTCACTATAGCG |
| T7 Reverse | TAGTTATTGCTCAGCGGTGG |
