## Supplementary figure for "A tau class Glutathione-S-Transferase (OsGSTU5) acts as a negative regulator of VirE2 interaction with T-DNA during *Agrobacterium* infection in rice"

| List of Y2H identified clones |
| --- |
| **1. LOC_Os04g36620** |
| **2. LOC_Os09g20220** |
| **3. LOC_Os04g48730** |
| **5. LOC_Os06g51220** |
| **6. LOC_Os02g52290** |
| **7. LOC_Os02g55890** |
| **8. LOC_Os08g42030** |
| **9. LOC_Os05g07030** |

Figure S1- List of VirE2 interacting clones in rice.

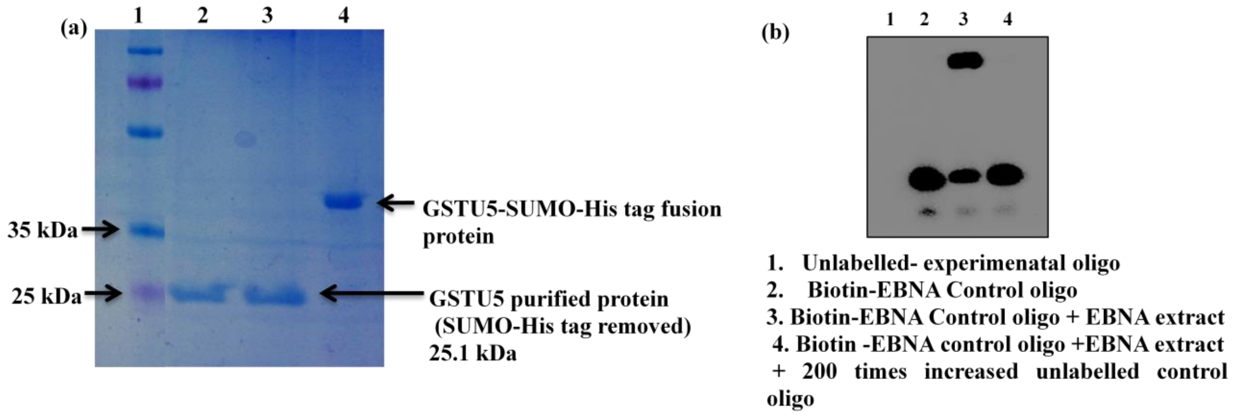


**Figure S2.**

**(a)** Coomassie stained SDS-PAGE showing GSTU5-SUMO fusion protein (38.1 kDa) (lane 4) and GSTU5 protein (25.1 kDa) (lane 2, 3) after removal of the his tag with SUMO protease **(b)** EMSA control reaction. EBNA stands for Epstein-Barr Nuclear Antigen. For control reaction 20 fmol of the biotin-control oligo and 1 unit of the EBNA extract were added in experimental mixture contining 1X binding buffer, 2.5% glycerol, 5 mM $MgCl_2$, 50 ng/µl poly (dI•dC), 0.05% NP-40).

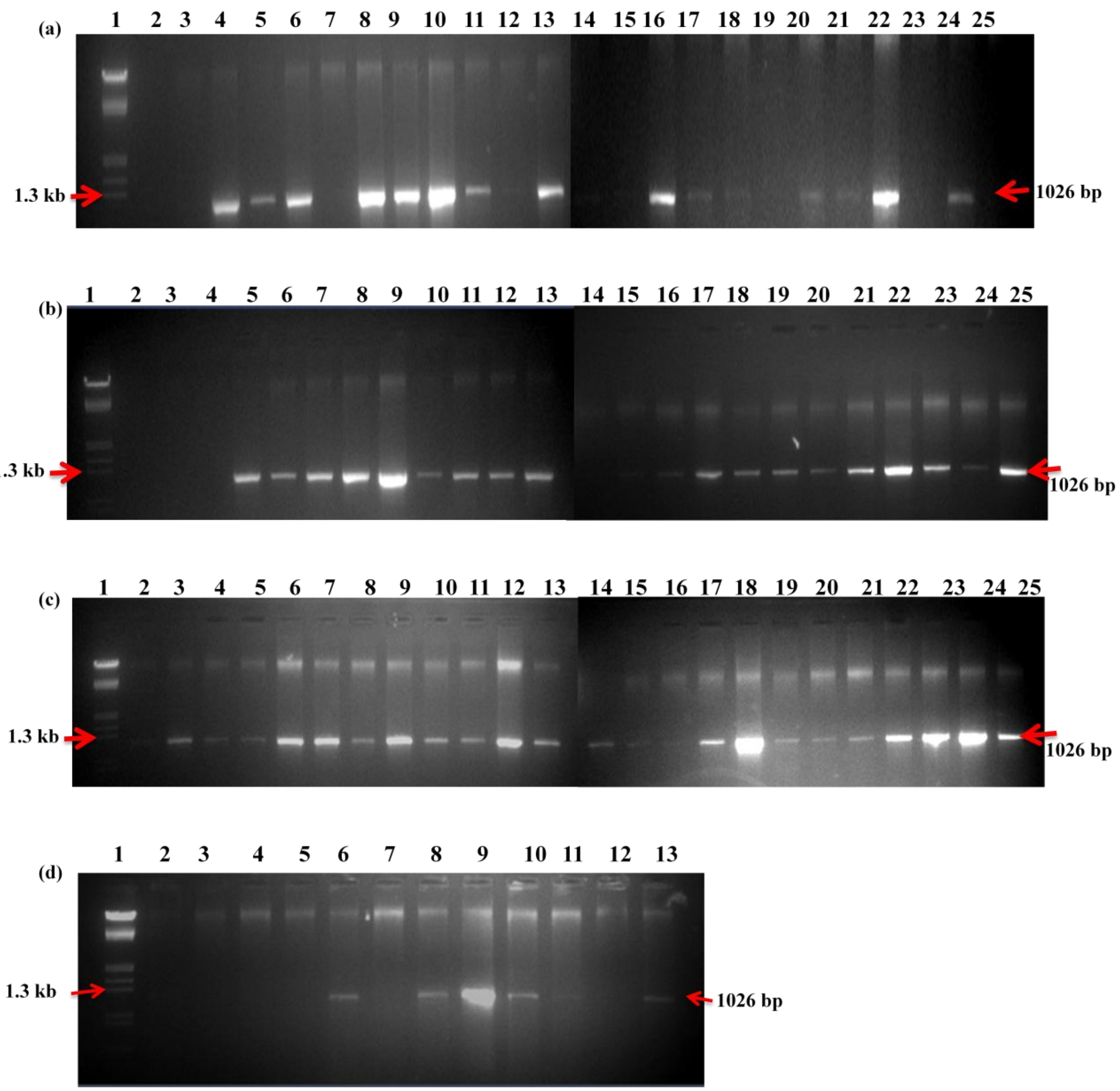


**Figure S3. Transformed rice plants and amplified with HPTII (Hygromycin) primers. The resulted PCR product ran on 1% Agarose gel.**

**(a)** Rice transgenic plants transformed with EV: lane 1 marker,2 negative control. Lane 3- 25, DNA samples. **(b and c)** Rice transgenic plants transformed with KD-GSTU5, lane marker, lane 2 negative control. **(b)** Lane 3-25 DNA samples. **(c)** Lane 2-25 DNA Samples. **(d)** Rice transgenic plants transformed with OE-GSTU5, (Overexpressed GSTU5). lane 1 marker, lane 2-13 DNA samples.

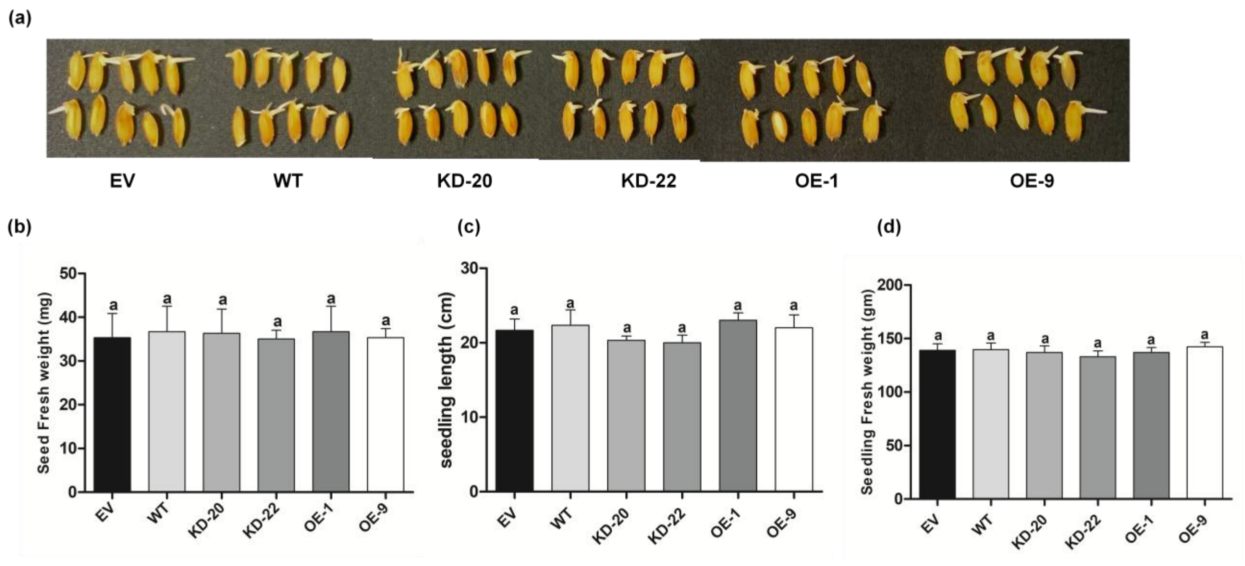


Figure S4. Comparison of WT and transgenic rice.

(a) Seed germination (b) seed weight (c) seedling length (d) seedling fresh weight.
